# Diet-Seg: Dynamic Hardness-Aware Learning for Enhanced Brain Tumor Segmentation

**DOI:** 10.1101/2025.05.31.657149

**Authors:** Boya Lv, Xinhang Huang, Qi Zhou, Mingxuan Li, Xinqing Xiao, Fuyi Li

## Abstract

Accurate brain tumor segmentation in magnetic resonance imaging (MRI) remains a critical challenge due to complex tumor heterogeneity, fuzzy boundaries, and significant inter-patient variability. In this study, we propose **Diet-Seg** (Difficulty-Informed Edge-enhanced Tiny Segmentation), a novel segmentation framework that integrates entropy-based pixel-wise hardness estimation into the training process via a dynamic learning rate modulation strategy. Specifically, we employ a pretrained 3D U-Net information model to quantify voxel-level prediction uncertainty, which is then used to guide the optimization of the main segmentation model. Diet-Seg is further enhanced by an RWKV-based U-Net backbone to capture global spatial dependencies and an EdgeNet module to preserve tumor boundaries through edge-aware fusion. Extensive experiments on the BraTS2018–2021 datasets demonstrate that Diet-Seg consistently outperforms state-of-the-art baselines across all tumor subregions. Notably, Diet-Seg achieves superior generalization when trained on one dataset and validated across multiple years. Moreover, the hardness maps offer interpretable insights into segmentation difficulty, potentially enabling human-AI collaboration in clinical practice. These results highlight the promise of entropy-aware training as a general strategy for robust and efficient medical image segmentation. The work is implemented in the open-source project available on GitHub (https://github.com/ManuelTurner/Diet-Seg)

## 1 Introduction

Gliomas, the most common type of malignant primary brain tumors[1], present significant diagnostic and therapeutic challenges due to their heterogeneous morphology and infiltrative growth[2]. Accurate segmentation of gliomas from magnetic resonance imaging (MRI) plays a critical role in pre-surgical planning, radiotherapy guidance, and prognosis prediction [3]. Multimodal MRI—including T1, T1-contrast, T2, and FLAIR sequences—provides complementary information that is essential for delineating tumor subregions[4], such as the enhancing tumor (ET), tumor core (TC), and whole tumor (WT)[5][6]. Nevertheless, the diversity in appearance across patients and modalities renders precise and robust subregion segmentation a non-trivial task.

Recent advances in deep learning, particularly convolutional neural networks (CNN) and transformer-based architectures[7][8], have markedly advanced the performance of brain tumor segmentation tasks [9, 10, 11]. Among these, 3D U-Net variants incorporating deep supervision and self-ensembling strategies, such as the model proposed by Théophraste Henry *et al*. (a top-10 solution in the BraTS 2020 challenge) [12], have established strong baselines within the BraTS benchmark. While these models effectively integrate volumetric context, their reliance on convolutional operations limits their capacity to capture long-range spatial dependencies.

To address these limitations, transformer-based methods have been introduced, offering global receptive fields and richer contextual reasoning capabilities[13]. However, their quadratic computational complexity in spatial dimensions poses significant challenges, especially in volumetric medical imaging[14].

Recent hybrid architectures, such as RWKV-UNet, effectively combine the efficiency of CNN with the global modeling capacity of Receptance Weighted Key Value (RWKV) modules [15]. RWKV-UNet has demonstrated state-of-the-art segmentation performance across multiple benchmarks while significantly reducing both parameter count and memory usage. Inspired by this, we investigate a lightweight yet powerful RWKV-based backbone for medical image segmentation.

Despite these architectural advances, another critical aspect remains underexplored: the structured uncertainty and varying learning difficulty across different tumor regions [16]. Most existing methods typically optimize all regions equally, neglecting the fact that tumor boundaries and infiltrative zones are inherently more ambiguous and challenging to learn [17]. Drawing inspiration from PH-Net, which introduced patchwise hardness estimation for breast lesion segmentation, we adopt an entropy-based uncertainty metric to quantify local hardness[18]. This metric captures per-patch confusion using normalized Shannon entropy over class probabilities, derived from an initial foundation model trained on BraTS2019 using the framework of Isensee *et al*.

Beyond capturing semantic features, accurate boundary delineation is vital for clinical applicability [19]. Motivated by edge-aware designs in multimodal segmentation networks, we introduce a dedicated edge detection module to enhance fine-grained localization [20]. This module incorporates CNN-based edge enhancement mechanisms to refine boundary precision, particularly in regions where semantic segmentation alone proves inadequate [21]. While prior studies have proposed multi-branch architectures for feature fusion, our approach integrates edge and semantic features in a more unified manner, enabling the model to jointly reason about anatomical shape and class identity.

We propose Diet-Seg (Difficulty-Informed Edge-enhanced Tiny Segmentation), a unified framework that incorporates three core components: (1) a lightweight RWKV-based backbone for efficient and expressive feature representation; (2) an EdgeNet module that captures and refines structural boundary details; and (3) a hardness-guided learning rate modulation scheme, where patches with higher entropy (more difficult regions) are trained with lower learning rates to ensure stable convergence and detail preservation, while easier regions benefit from faster learning. This strategy acts as a fine-grained curriculum, dynamically adjusting the optimization difficulty in space and time.

Empirical evaluation on the BraTS2020 validation dataset demonstrates that our proposed Diet-Seg consistently outperforms the original foundation model. Specifically, the mean Dice scores across tumor subregions improved from ET: 84.81%, TC: 81.79%, and WT: 79.82% to ET: 86.53%, TC: 86.86%, and WT: 81.91%. The most significant improvements were observed in the segmentation of the enhancing entire tumor and tumor core regions. Importantly, Diet-Seg achieves these performance gains with only half the number of parameters compared to the original 3D U-Net teacher model, underscoring both the efficacy and computational efficiency of our proposed framework.

## 2 Methods

This section outlines the key components of our proposed framework, Diet-Seg. These include an entropy-based hardness estimation module, an RWKV-UNet backbone for both global and local feature encoding, and an edge-enhancement module designed to improve structural boundary delineation.

### 2.1 Hardness Estimation Module

To adaptively guide the training process based on spatial prediction uncertainty, we propose an entropy-based *hardness estimation module*. This module leverages the entropy of class probability distributions to quantify local uncertainty, producing patch-wise hardness maps that dynamically reflect regional complexity and confusion in the segmentation predictions.

#### 2.1.1 Motivation

Conventional segmentation approaches uniformly weigh all pixels during training, regardless of regional uncertainty[22]. However, in medical imaging—especially brain tumor segmentation—boundary and heterogeneous tissue regions exhibit significantly higher uncertainty[23]. By explicitly quantifying and 4 Diet-Seg modeling these spatial variations, the training procedure can prioritize challenging regions, leading to more precise and stable optimization.

#### 2.1.2 Class Probability Correction

The foundation segmentation model outputs overlapping or cumulative probability maps due to hierarchical tumor structures (whole tumor, tumor core, enhancing tumor). Mutually exclusive probability distributions are required to compute entropy accurately. Let the model output probability tensor be defined as:

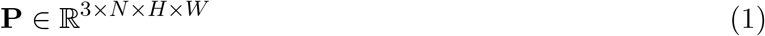

with channels corresponding to enhancing tumor (ET, **P**_0_), tumor core (TC, **P**_1_), and whole tumor (WT, **P**_2_). We perform the following correction to obtain non-overlapping class probabilities 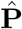 :

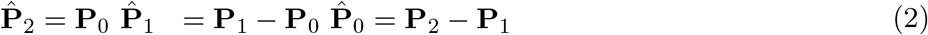

These corrected probabilities are then clipped between [0, 1] and normalized to maintain numerical stability:

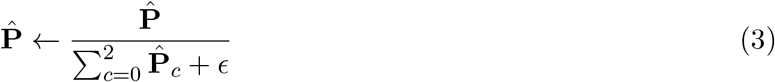

where *ϵ* (e.g., 10^−5^) prevents division by zero and ensures numerical robustness.

#### 2.1.3 Patch-wise Entropy Calculation

To quantify local uncertainty, the normalized entropy is computed for each non-overlapping spatial patch within each slice. We partition each slice into patches of size *s* × *s*, and for each patch *m*, we calculate the entropy-based hardness score *r*_*m*_ as:

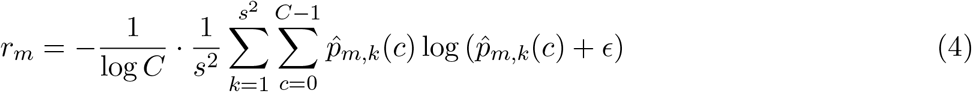

Where 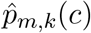 is the corrected probability for class *c* at pixel *k* within patch *m, C* = 3 represents the number of segmentation classes, and the normalization factor 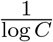 ensures that *r*_*m*_ is bounded between 0 and 1.

This formulation implies that patches with high prediction uncertainty (uniform probability distributions) yield higher entropy, indicating increased learning complexity.

#### 2.1.4 Slice-wise Hardness Normalization

Since medical images consist of multiple slices with varying levels of uncertainty, we normalize the hardness scores within each slice *i* independently to maintain comparability across different spatial contexts:

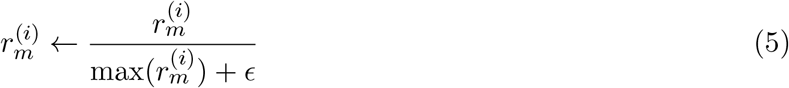

This normalization prevents individual slices with inherently higher uncertainty from disproportionately influencing the training dynamics.

#### 2.1.5 Hardness Map Upsampling

Patch-level hardness values are upsampled back to the original spatial resolution to apply hardness scores at pixel-level granularity. Each pixel within a patch inherits the hardness score of that patch:

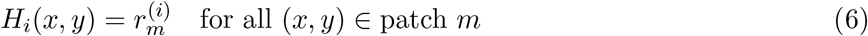

The resulting per-pixel hardness map *H* ∈ R^*N* ×*H*×*W*^ visually and numerically highlights regions with varying degrees of prediction difficulty, serving as a direct guide for adaptive training and optimization.

#### 2.1.6 Practical Implications

The entropy-based hardness map not only facilitates adaptive loss weighting, where harder regions (typically boundaries or heterogeneous tissues) receive more careful gradient updates, but also acts as an interpretability tool, clearly visualizing areas of uncertainty in predictions. This approach ultimately enhances model accuracy and robustness, particularly in challenging medical segmentation tasks.

### 2.2 RWKV-UNet Backbone

Modeling both local and long-range dependencies in medical image segmentation is critical for accurately delineating tumor boundaries and understanding structural context[24]. To this end, we develop a lightweight yet expressive encoder-decoder architecture, termed **RWKV-UNet**, which integrates the Receptance Weighted Key Value (RWKV) mechanism into a hierarchical U-Net design.

#### 2.2.1 Motivation

CNN are widely used in segmentation networks due to their strong local feature extraction capability[25]. However, they are inherently limited in modeling long-range dependencies due to restricted receptive fields[10]. While transformer-based architectures provide global attention, they often incur quadratic computational complexity with respect to image resolution, making them suboptimal for dense prediction tasks[26].

The RWKV architecture, originally proposed for efficient sequence modeling, combines the global receptive field of attention with the computational efficiency of recurrent processing. We adapt and extend RWKV to spatial domains, enabling our model to achieve rich contextual understanding without the high memory footprint of traditional transformers.

#### 2.2.2 Architecture Overview

The RWKV-UNet follows a standard U-Net topology consisting of a multi-stage encoder, a feature fusion bottleneck, and a symmetric decoder. Each encoder block is implemented using an `iR_RWKV` module, which includes:

- Pointwise convolutions for feature projection and expansion.
- RWKV-based attention for capturing long-range spatial context.
- Channel-wise squeeze-and-excitation (SE) layers for recalibration.
- Depthwise convolutions for local detail refinement.

Decoder blocks use `UpBlock` modules that upsample feature maps and integrate skip connections. A Cross-Channel Mix (CCM) module is used at the bottleneck to aggregate multi-scale encoder features.

#### 2.2.3 RWKV Spatial Attention

Let the flattened spatial input be **X** ∈ R^*B*×*T* ×*C*^, where *B* is the batch size, *T* = *H* × *W* is the spatial dimension, and *C* is the number of channels. The spatial RWKV attention mechanism computes the output **Y** ∈ R^*B*×*T* ×*C*^ as:

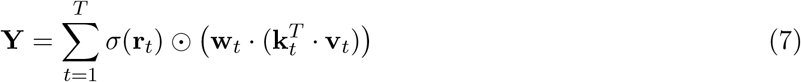

where **k**_*t*_ and **v**_*t*_ are the key and value embeddings, **r**_*t*_ is the receptance gate, and **w**_*t*_ represents a learnable decay factor.

To enhance spatial generalization, we perform a spatial channel shift operation before key-value projection. Let **X**^′^ denote the shifted feature map and *α* be the learned mixing coefficient, the final input becomes:

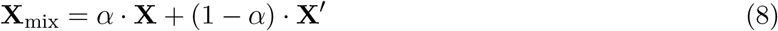

The resulting **X**_mix_ is then passed through separate linear layers to obtain **K, V, R** and processed by the CUDA-optimized RWKV kernel.

#### 2.2.4 Cross-Channel Mixing and Decoder Design

The decoder consists of symmetric upsampling blocks, each performing bilinear upsampling followed by convolution, SE, and normalization. A CCM module at the bottleneck fuses high-level encoder features from different resolutions:

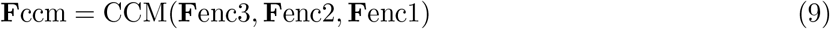

This design ensures both fine and coarse semantic features are preserved, resulting in better delineation of tumor subregions.

### 2.3 EdgeNet: Boundary-Aware Feature Enhancement

Accurate segmentation in medical images not only requires correct classification of regions but also precise delineation of object boundaries—particularly for tasks such as brain tumor subregion segmentation, where subtle differences in tissue interfaces are clinically significant. To address this, we introduce **EdgeNet**, a lightweight boundary-aware enhancement module that augments semantic features with high-frequency edge information.

#### 2.3.1 Motivation

Despite recent advances in segmentation networks, existing models often struggle with capturing sharp transitions and boundaries, especially in low-contrast MRI modalities[27]. Typical decoder designs focus heavily on semantic restoration, overlooking spatial gradients that define anatomical boundaries[28]. EdgeNet explicitly models such spatial discontinuities, thereby enforcing spatial consistency and improving region separation.

#### 2.3.2 Architecture Overview

EdgeNet operates in parallel with the main decoder stream and enhances intermediate features with boundary-specific signals. As shown in Figure **??**, low-level encoder features **F**_*l*_ at different stages are first projected to a lower-dimensional space using 1 × 1 convolutions:

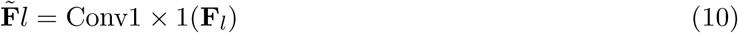

where *l* indexes the encoder level. The projected features are then passed to the edge detection unit.

#### 2.3.3 Boundary Extraction via Gradient Enhancement

To emphasize spatial discontinuities, we compute the residual between original and smoothed features via local averaging:

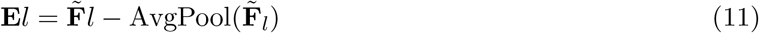

where **E**_*l*_ represents the edge-enhanced map, and the average pooling is implemented with a 3×3 kernel and stride 1. This operation highlights high-frequency spatial variations, which typically correspond to object boundaries.

The resulting edge features are then fused with semantic representations in the cross-channel mixing block (CCM), enabling joint reasoning over both region-wise context and local structure:

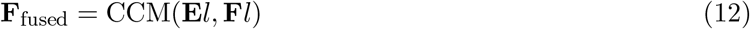

#### 2.3.4 Integration and Training

EdgeNet modules are deployed at multiple encoder depths and contribute to the fusion pathway between encoder and decoder. Importantly, this enhancement does not introduce significant overhead as it relies on lightweight convolution and pooling operations. During training, the edge-aware pathways act as implicit regularizers, guiding the backbone to maintain sharp boundaries across tumor subregions.

By leveraging explicit boundary modeling alongside RWKV-based semantic encoding, EdgeNet helps the network achieve finer structural prediction and improved generalization across challenging cases.

### 2.4 Loss Function

To effectively optimize the segmentation model under complex spatial variations, we formulate a composite loss function that integrates pixel-wise cross-entropy, Dice similarity loss, and a background confidence regularization term. Central to our formulation is a hardness-aware weighting mechanism that adaptively emphasizes ambiguous or difficult regions during training.

#### 2.4.1 Overall Formulation

The total loss is expressed as:

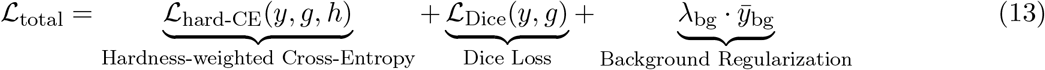

where *y* ∈ R^*C*×*H*×*W*^ denotes the predicted probability map, *g* is the ground truth label map, *h* is the per-pixel hardness map as described in Section **??**, and *λ*_bg_ is a weighting coefficient for background regularization, empirically set to 1.

#### 2.4.2 Hardness-Weighted Cross-Entropy Loss

To guide the network toward uncertain or error-prone regions, we employ a spatial weighting strategy based on the hardness map. The hardness-weighted cross-entropy is defined as:

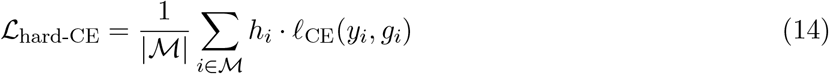

where *l*_CE_(*y*_*i*_, *g*_*i*_) denotes the standard cross-entropy loss at pixel *i, h*_*i*_ ∈ [0, 1] is the corresponding hardness weight, and M represents the set of valid pixels excluding ignored labels. The hardness map *h* is detached during backpropagation to prevent gradient interference.

#### 2.4.3 Dice Loss

To directly optimize for spatial overlap and address class imbalance, we incorporate the Dice loss:

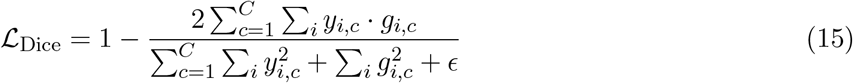

where *y*_*i,c*_ and *g*_*i,c*_ denote the predicted and ground truth one-hot labels for class *c*, and *ϵ* is a small constant to ensure numerical stability.

#### 2.4.4 Background Confidence Regularization

Instead of assuming an explicit background class, we define background pixels as those where the model exhibits uniformly low confidence across all foreground classes. Specifically, a pixel is considered background if the predicted logits for all classes are negative:

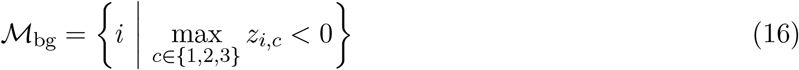

where *z*_*i,c*_ denotes the raw logit for class *c* at pixel *i*.

#### 2.4.5 Optimization Strategy

The total loss is minimized via stochastic gradient descent. All terms are jointly optimized in an end-to-end manner. The proposed loss formulation not only improves prediction fidelity in structurally complex and uncertain regions but also enhances generalization by explicitly encoding prediction difficulty into the learning objective.

## 3 Experiments

### 3.1 Datasets

We evaluate the proposed framework on two publicly available brain tumor segmentation datasets: **BraTS2019** and **BraTS2020** [29]. BraTS2019 contains 260 subjects, and BraTS2020 comprises 369 subjects. Each sample includes four MRI modalities—T1, T1ce, T2, and FLAIR—with a spatial resolution of 240 × 240 × 155. Ground-truth annotations consist of three tumor subregions: ET, TC, and WT. We employ 5-fold cross-validation to ensure robust evaluation.

### 3.2 Experimental Setup

**Preprocessing and Sampling** All MRI volumes are preprocessed by standardizing intensity values and cropping to the non-zero region of interest. During training, we randomly sample sub-volumes of size 128 × 128 × 128 from each case to ensure spatial diversity.

**Foundation Model for Hardness Estimation** To compute pixel-wise uncertainty (hardness), we train a 3D U-Net model on BraTS2019 following the setup of a top-10 ranked solution from the BraTS2019 challenge. This model, referred to as the *information model*, serves as the backbone for entropy-based hardness estimation. It is trained for 400 epochs using the Adam optimizer and a fixed learning rate schedule. The same model is validated on the BraTS2018, BraTS2019, BraTS2020, and BraTS2021 datasets to ensure generalization.

**Diet-Seg Model Training** Our proposed segmentation framework, *Diet-Seg*, is trained on both BraTS2019 and BraTS2020 using a dynamic hardness-aware learning rate schedule. Specifically, the base learning rate is adaptively modulated by patch-wise entropy values derived from the foundation model. Learning rate annealing is performed every 400 iterations, with a total of 2000 training epochs. The model is optimised using the Adam optimizer with an initial learning rate of 0.001 and a batch size of 3.

**Implementation Details** All models are implemented in PyTorch and trained on a single NVIDIA RTX 4090 GPU. To accelerate RWKV inference, we compile the CUDA-based kernel with C++ using custom wrappers for optimal GPU utilization. The default image size during inference is 128 × 128 × 128. For reproducibility, random seeds are fixed to 1234.

### 3.3 Quantitative Analysis

As shown in Tables 1 and 2, our proposed model, **Diet-Seg**, achieves superior performance on both the BraTS2020 and BraTS2019 datasets in terms of Dice Similarity Coefficient (DSC) and 95% Hausdorff Distance (HD95). Compared to multiple state-of-the-art (SOTA) segmentation methods, Diet-Seg yields consistently better results across all tumor subregions: enhancing tumor (ET), whole tumor (WT), and tumor core (TC).

**Table 1:**
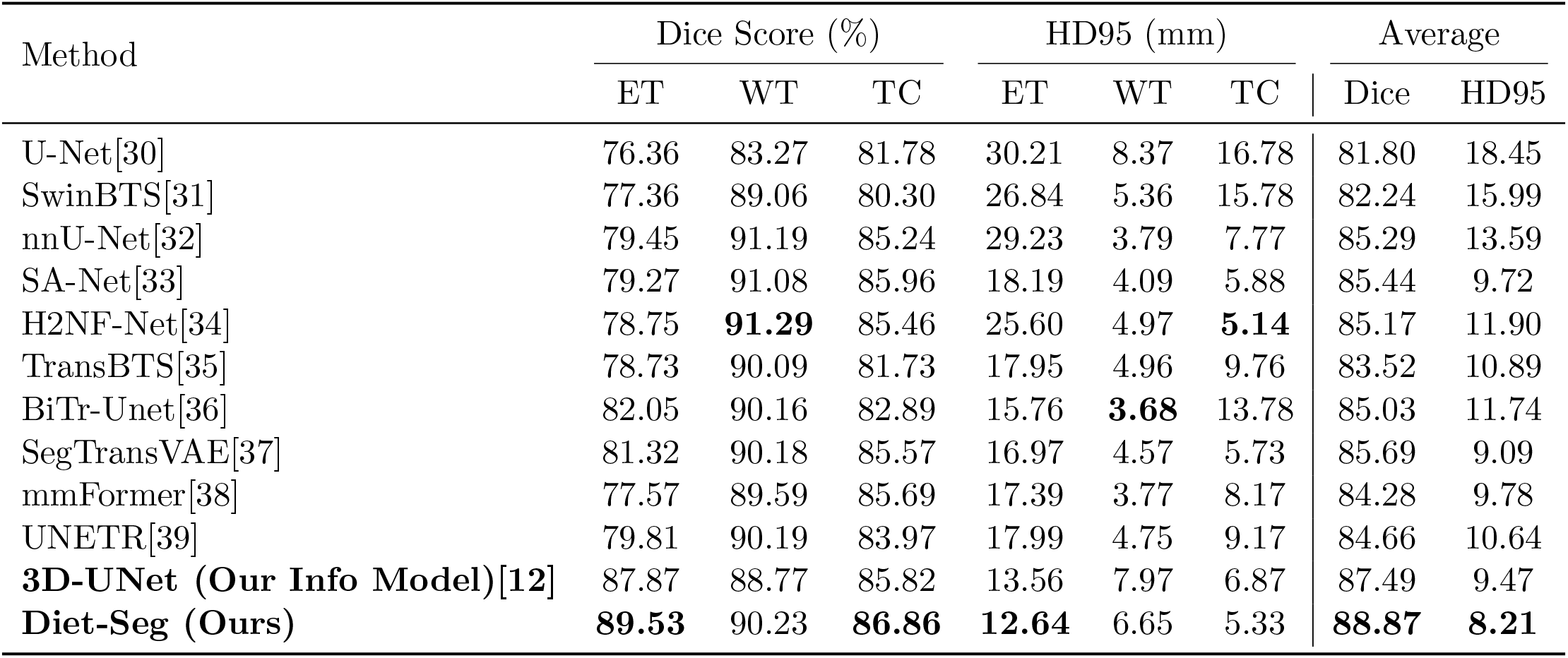
Quantitative comparison of segmentation performance (Dice Score & HD95) on the BraTS2020 dataset. The best-performing results among baseline and proposed methods are highlighted in **bold**.

| Method | Dice Score (%) |  |  | HD95 (mm) |  |  | Average |  |
| --- | --- | --- | --- | --- | --- | --- | --- | --- |
|  | ET | WT | TC | ET | WT | TC | Dice | HD95 |
| U-Net[30] | 76.36 | 83.27 | 81.78 | 30.21 | 8.37 | 16.78 | 81.80 | 18.45 |
| SwinBTS[31] | 77.36 | 89.06 | 80.30 | 26.84 | 5.36 | 15.78 | 82.24 | 15.99 |
| nnU-Net[32] | 79.45 | 91.19 | 85.24 | 29.23 | 3.79 | 7.77 | 85.29 | 13.59 |
| SA-Net[33] | 79.27 | 91.08 | 85.96 | 18.19 | 4.09 | 5.88 | 85.44 | 9.72 |
| H2NF-Net[34] | 78.75 | **91.29** | 85.46 | 25.60 | 4.97 | **5.14** | 85.17 | 11.90 |
| TransBTS[35] | 78.73 | 90.09 | 81.73 | 17.95 | 4.96 | 9.76 | 83.52 | 10.89 |
| BiTr-Unet[36] | 82.05 | 90.16 | 82.89 | 15.76 | **3.68** | 13.78 | 85.03 | 11.74 |
| SegTransVAE[37] | 81.32 | 90.18 | 85.57 | 16.97 | 4.57 | 5.73 | 85.69 | 9.09 |
| mmFormer[38] | 77.57 | 89.59 | 85.69 | 17.39 | 3.77 | 8.17 | 84.28 | 9.78 |
| UNETR[39] | 79.81 | 90.19 | 83.97 | 17.99 | 4.75 | 9.17 | 84.66 | 10.64 |
| **3D-UNet (Our Info Model)[12]** | 87.87 | 88.77 | 85.82 | 13.56 | 7.97 | 6.87 | 87.49 | 9.47 |
| **Diet-Seg (Ours)** | **89.53** | 90.23 | **86.86** | **12.64** | 6.65 | 5.33 | **88.87** | **8.21** |

**Table 2:** Quantitative comparison of segmentation performance (Dice Score & HD95) on the BraTS2019 dataset. The best results are highlighted in **bold**.

| Method | DSC (%) |  |  | HD95 (mm) |  |  | Average |  |
| --- | --- | --- | --- | --- | --- | --- | --- | --- |
|  | ET | WT | TC | ET | WT | TC | DSC | HD95 |
| U-Net[30] | 73.34 | 88.26 | 75.98 | 6.21 | 6.17 | 8.68 | 79.19 | 7.02 |
| Attention U-Net[40] | 75.96 | 88.81 | 77.20 | 5.20 | 7.76 | 8.26 | 80.65 | 7.07 |
| SA-LuT-Net [41] | 78.21 | 90.34 | 84.82 | **3.69** | 4.46 | 5.36 | 84.60 | 4.47 |
| TransBTS | 78.90 | 90.01 | 81.90 | 4.75 | 4.64 | 6.04 | 83.60 | 5.47 |
| BiTr-Unet | 81.09 | 90.46 | 81.78 | 4.61 | 4.73 | 7.82 | 84.11 | 6.06 |
| SegTransVAE | 80.85 | 90.15 | 85.57 | 4.14 | 4.47 | 5.67 | 85.41 | 5.09 |
| UNETR | 80.23 | 90.27 | 84.91 | 5.23 | 4.53 | 6.52 | 85.14 | 5.43 |
| **3D-UNet (Our Info Model)** | 84.81 | 85.82 | 81.79 | 6.26 | 12.31 | 7.59 | 84.14 | 8.72 |
| **Diet-Seg (Ours)** | **86.53** | **89.91** | **86.86** | 5.21 | **8.97** | **5.33** | **87.77** | **6.50** |

**Table I — Comparison on BraTS2020** Table 1 shows the comparative performance of Diet-Seg against 12 representative methods on the BraTS2020 dataset, including U-Net, SwinBTS, nnU-Net, SA-Net, H2NF-Net, TransBTS, BiTr-Unet, SegTransVAE, mmFormer, and UNETR. Among these, our **Diet-Seg** model achieves the best performance with Dice scores of **89.53%**, 90.23%, and **86.86%** for ET, WT, and TC, respectively. The corresponding HD95 values are **12.64 mm**, 6.65 mm, and **5.33 mm**, yielding the best overall average Dice of **88.87%** and average HD95 of **8.21 mm**.

Compared to conventional CNN-based models like U-Net and nnU-Net, which attain 81.80% and 85.29% average Dice respectively, Diet-Seg demonstrates a significant improvement of 7.07% and 3.58%, respectively. Diet-Seg also outperforms transformer-based methods such as TransBTS (83.52% Dice, 10.89 mm HD95) and UNETR (84.66% Dice, 10.64 mm HD95), highlighting its ability to effectively model both long-range dependencies and local structural features.

It is also noteworthy that our baseline model, **3D-UNet (Info Model)**, which is used for hardness estimation, achieves a strong performance of 87.49% Dice and 9.47 mm HD95. Nevertheless, Diet-Seg improves upon this by an additional +1.38% in Dice and −1.26 mm in HD95. This indicates that while the Info Model is a powerful backbone for hardness-aware learning, it is through Diet-Seg’s dynamic curriculum strategy and edge-aware architecture that maximum segmentation effectiveness is achieved.

**Table II — Comparison on BraTS2019** Table 2 illustrates the results on the BraTS2019 dataset. Our **Diet-Seg** achieves Dice scores of **86.53%** (ET), **89.91%** (WT), and **86.86%** (TC), with HD95 values of **5.21 mm, 8.97 mm**, and **5.33 mm**, respectively. The overall average Dice and HD95 are **87.77%** and **6.50 mm**, respectively—again outperforming all other competing methods.

Among the baseline methods, SA-LuT-Net achieves competitive HD95 (4.47 mm) but lower DSC (84.60%), while SegTransVAE achieves 85.41% Dice and 5.09 mm HD95. Compared to these, Diet-Seg provides both higher accuracy and finer boundary segmentation. Moreover, compared to our own **3D-UNet Info Model**, which achieves 84.14% Dice and 8.72 mm HD95, Diet-Seg shows a 3.63% improvement in Dice and a 2.22 mm reduction in HD95.

**Table III — Cross-Year Generalization Analysis** Table 3 presents a comprehensive cross-year evaluation, where all models are trained exclusively on BraTS2019 and validated across BraTS2018, BraTS2019, BraTS2020, and BraTS2021. This setup enables a fair assessment of the model’s generalization capability under distributional shifts.

**Table 3:** Cross-year validation results. Models are trained on BraTS2019 and evaluated on BraTS2018, 2019, 2020, and 2021 datasets. All values represent Dice Score (%). Improvements of **Diet-Seg** over the Info Model are indicated in parentheses.

| Dataset | Info Model |  |  | Diet-Seg (Improvement) |  |  |
| --- | --- | --- | --- | --- | --- | --- |
|  | ET | TC | WT | ET | TC | WT |
| BraTS2018 | 81.33 | 82.54 | 79.50 | **84.85** (+4.33%) | **84.82** (+2.76%) | **80.70** (+1.51%) |
| BraTS2019 | 84.81 | 85.82 | 81.79 | **86.53** (+2.03%) | **89.91** (+4.76%) | **86.86** (+6.21%) |
| BraTS2020 | 72.67 | 68.78 | 70.68 | **76.57** (+5.37%) | **74.45** (+8.24%) | **72.35** (+2.36%) |
| BraTS2021 | 78.79 | 76.39 | 68.00 | **83.19** (+5.59%) | **80.14** (+4.91%) | **77.20** (+13.53%) |

Our **Diet-Seg** consistently outperforms the **Info Model** across all datasets and tumor subregions. On the BraTS2018 dataset, Diet-Seg improves Dice scores by **4.33%** (ET), **2.76%** (TC), and **1.51%** (WT), respectively. This confirms its robustness to older data distributions.

On the in-domain BraTS2019 set, where both models were trained, Diet-Seg still achieves noticeable gains, especially a **6.21%** boost in WT Dice and a **4.76%** increase in TC Dice, indicating that the hardness-aware training strategy enhances performance even in familiar domains.

For the more challenging BraTS2020 and BraTS2021 datasets, which contain significant structural and scanner variability, the performance gap further widens. Diet-Seg surpasses Info Model by up to **5.37%** (ET) and **8.24%** (TC) on BraTS2020, and by **13.53%** (WT) and **5.59%** (ET) on BraTS2021.

These results highlight Diet-Seg’s ability to generalize better under domain shifts and its resilience to noisy or shifted label distributions.

Overall, the consistent improvements across years and tumor subregions clearly demonstrate the superior generalization capacity of Diet-Seg, particularly in scenarios involving inter-domain variability and unseen data.

**Summary** In summary, our proposed Diet-Seg framework consistently outperforms all baseline models across both the BraTS2019 and BraTS2020 datasets. It not only surpasses the performance of classic CNN-based architectures and state-of-the-art Transformer variants, but also achieves substantial improvements over our own information model in terms of segmentation accuracy and boundary delineation. These results demonstrate the robustness and effectiveness of Diet-Seg, highlighting the value of integrating hardness-aware learning and edge enhancement strategies in medical image segmentation.

### 3.4 Qualitative Analysis

Figure 5 presents a comprehensive visual comparison between the 3D-UNet information model and the proposed Diet-Seg framework, using representative cases from the BraTS dataset. Each row represents a single patient slice, while the columns sequentially depict: (1) the input T1ce modality MRI slice, (2) the ground truth segmentation label, (3) prediction result from the Info Model, (4) entropy heatmap from the Info Model, (5) Diet-Seg prediction, (6) Diet-Seg entropy map, and (7) the final aggregated hardness map derived from multi-class uncertainty.

**Figure 1:**
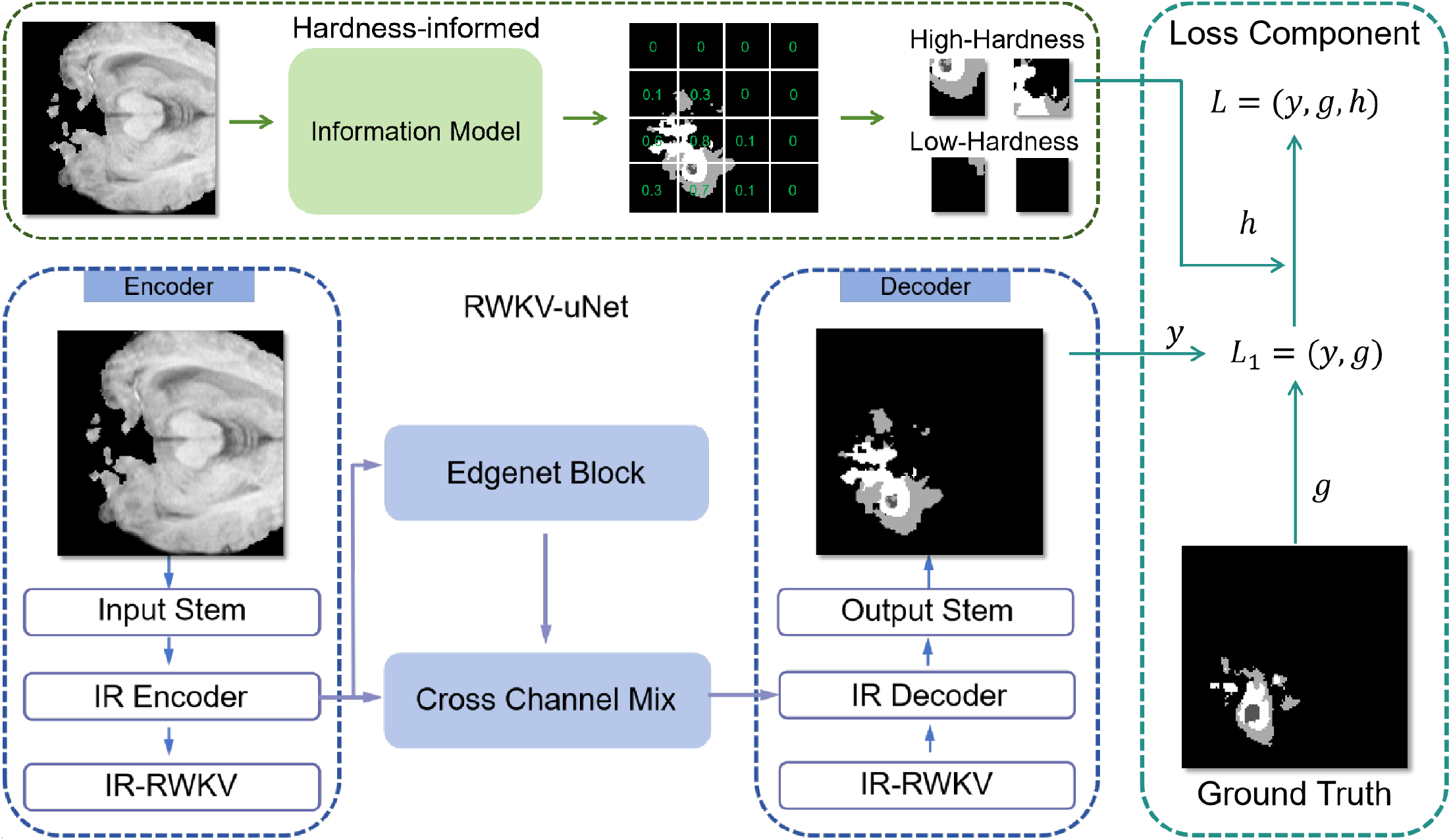
Overview of the Diet-Seg architecture. The proposed framework comprises three main components: an encoder based on RWKV blocks for hierarchical feature extraction, an EdgeNet module that enhances boundary representation, and a Cross-Channel Mix module that fuses semantic and edge features. The encoder processes multimodal MRI inputs and forwards features to both the decoder and the EdgeNet block. The EdgeNet explicitly extracts structural edge cues, which are combined with the semantic features through cross-channel interactions to refine spatial representations. The decoder then integrates multi-level features to generate accurate subregion segmentation masks. This architecture allows efficient modeling of both global context and fine-grained boundaries, while remaining lightweight.

**Figure 2:**
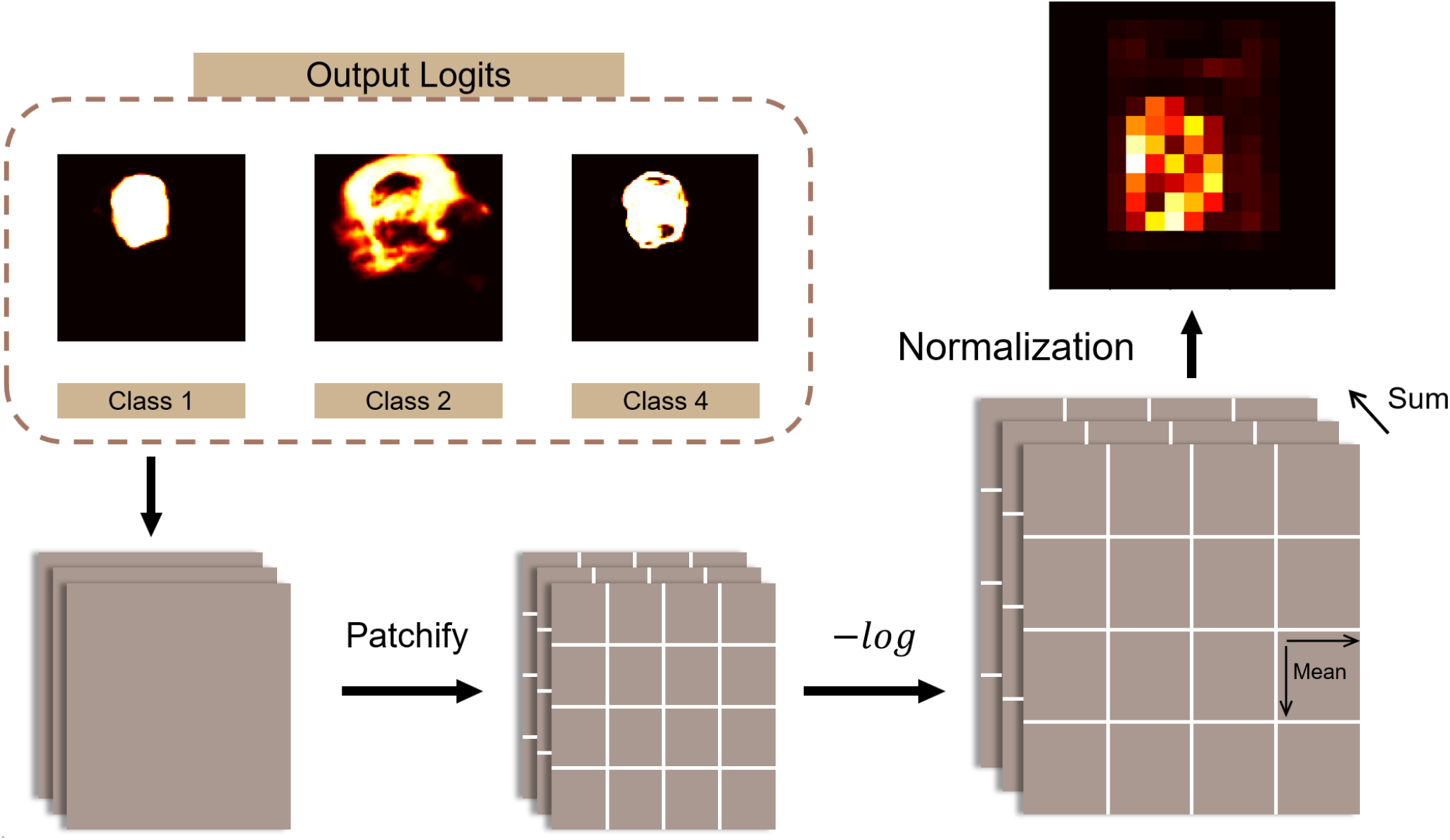
Pipeline of the entropy-based hardness estimation module.

**Figure 3:**
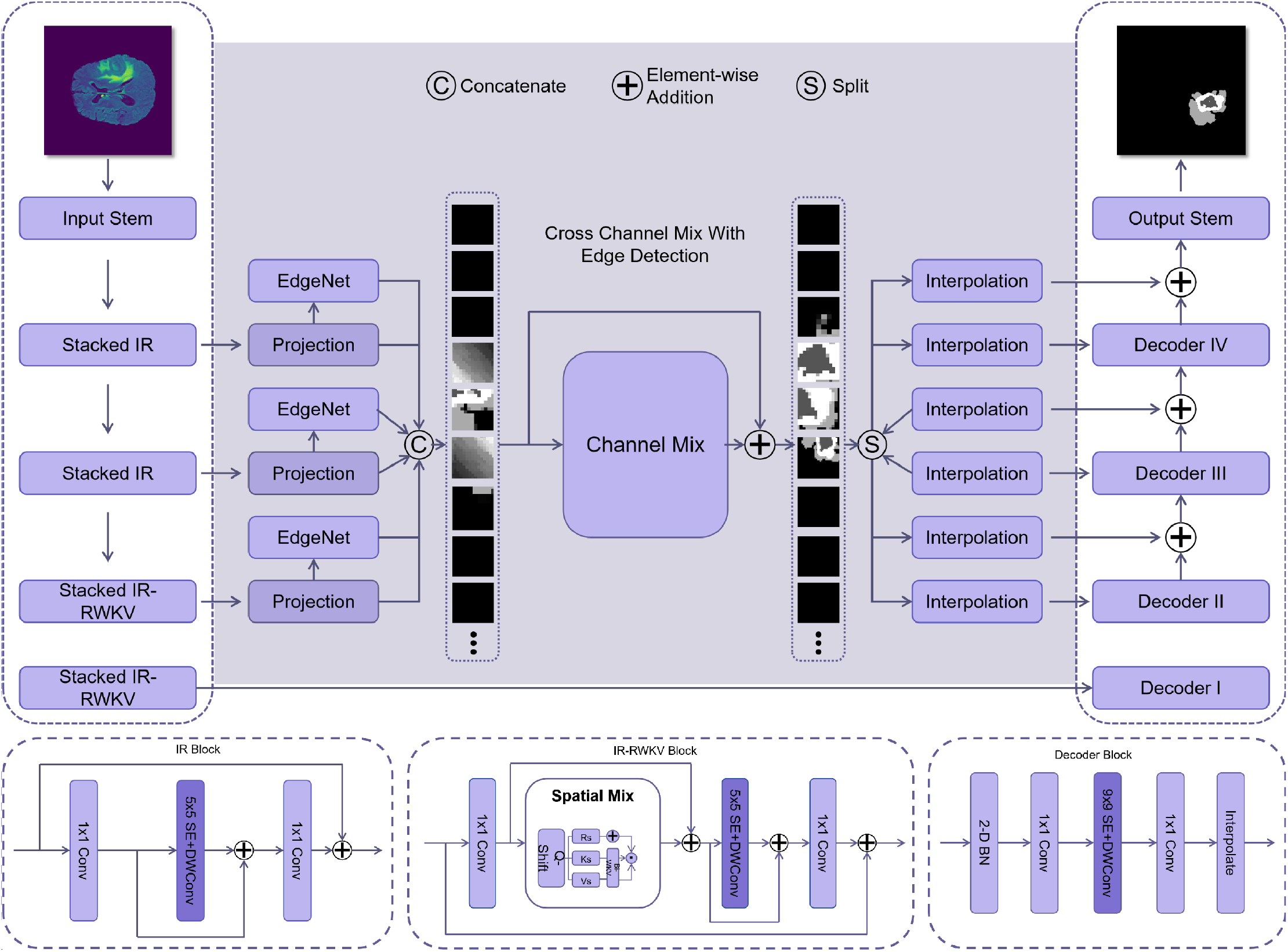
Overall architecture of the proposed RWKV-UNet framework.

**Figure 4:**
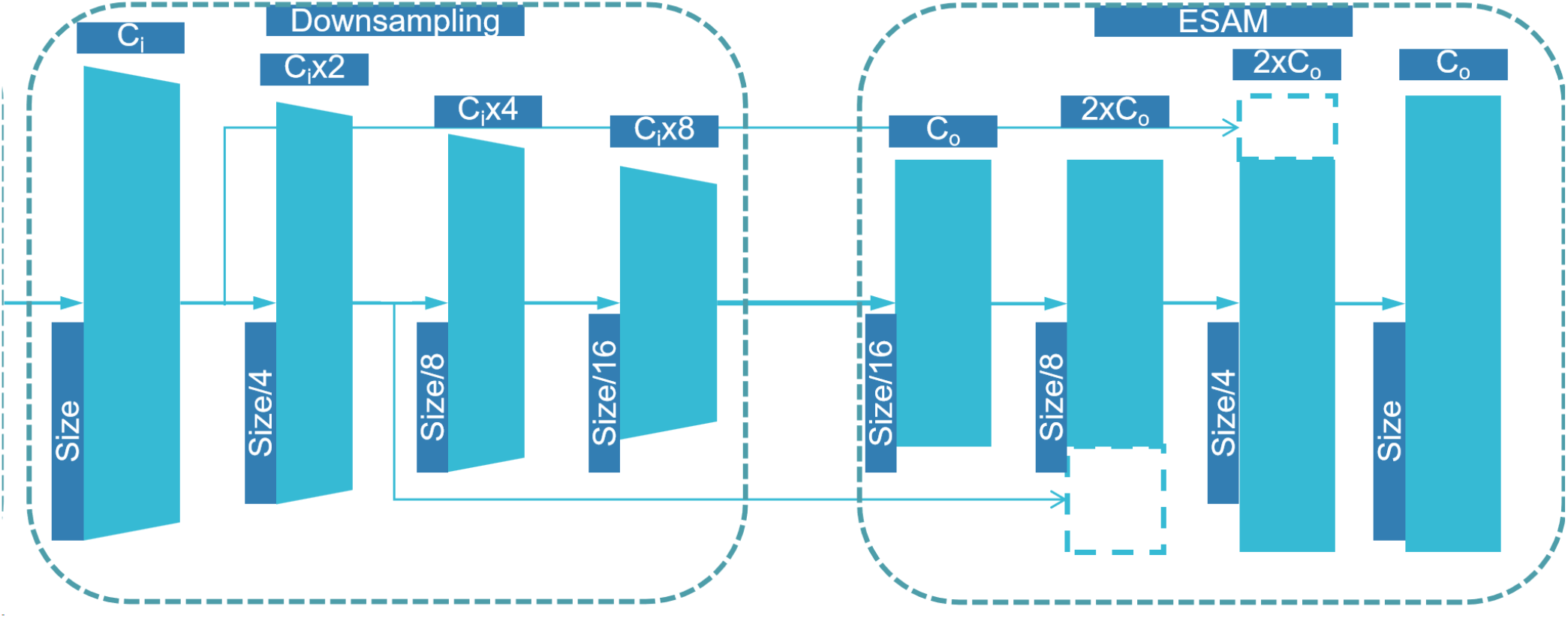
Illustration of the encoder-decoder structure with Edge-Aware Semantic Aggregation Module (ESAM).

**Figure 5:**
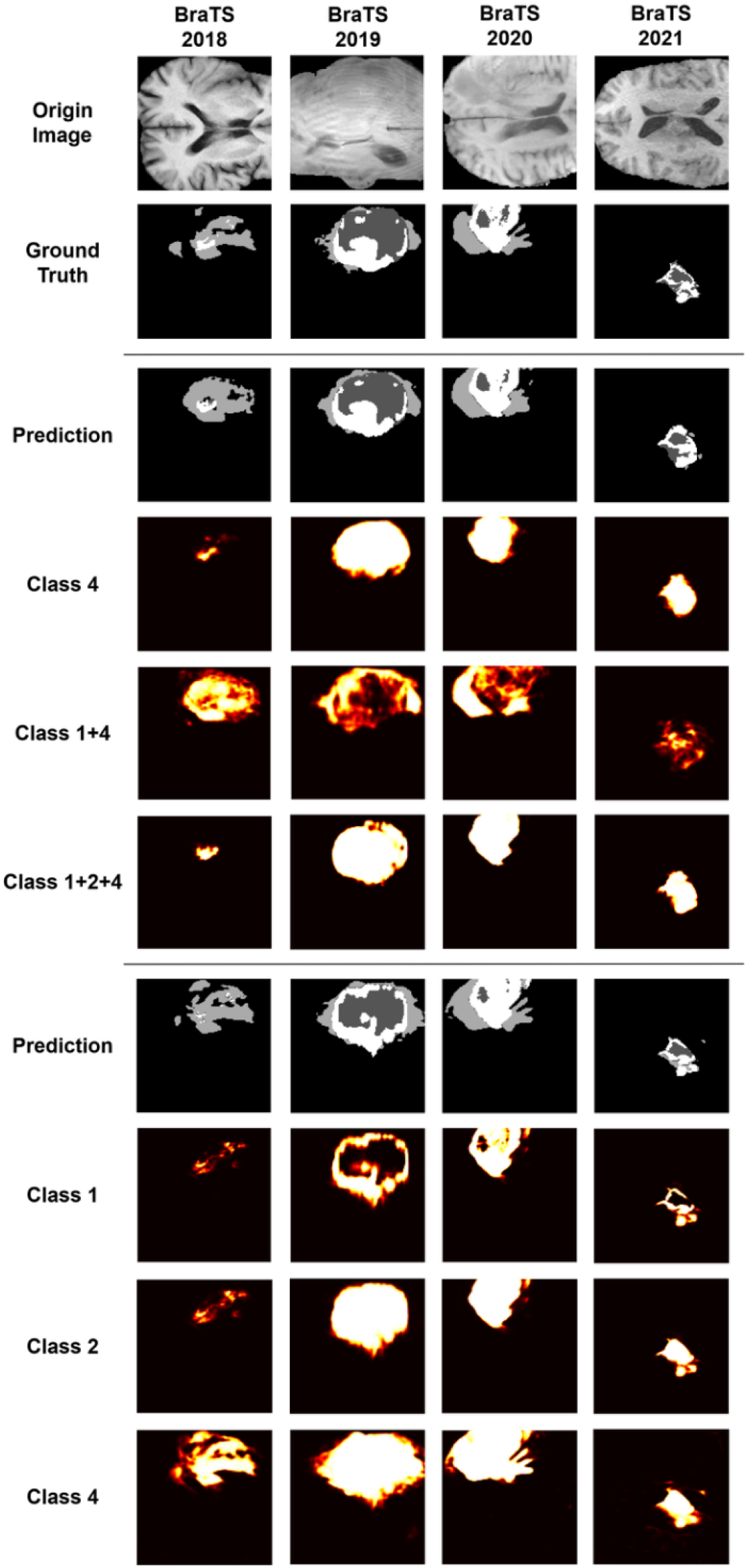
Qualitative comparisons of segmentation and uncertainty visualization.

From a structural perspective, Diet-Seg consistently generates more coherent and complete tumor segmentations. In several cases (*e.g*., row 2 and row 4), the Info Model shows fragmented or overly conservative delineation of tumor subregions, particularly the ET and TC, whereas Diet-Seg successfully identifies the correct tumor boundaries and captures the full extent of the lesion.

Edge sharpness is also significantly improved. The entropy maps of the Info Model are characterized by diffuse uncertainty across the tumor region and often spill over into healthy tissue, indicating less reliable boundary prediction. In contrast, Diet-Seg produces entropy distributions that are tightly localized to the lesion perimeter, revealing that the model is more confident inside tumor areas and more cautious at semantic boundaries—an expected and desirable behavior.

The last column shows the aggregated pixel-wise hardness map, computed from the mean entropy across all tumor classes. These maps visualize regions of high semantic ambiguity that challenge the segmentation network. We observe that high-hardness regions often coincide with tumor borders and small or irregularly shaped lesions. This suggests that hardness learning effectively identifies “difficult” regions during training and allows Diet-Seg to adapt its learning focus accordingly.

Additionally, we observe that Diet-Seg is more robust in cases with large tumor mass (e.g., row 3), where the Info Model fails to delineate the internal heterogeneity of the lesion. Conversely, for small lesions with limited contrast (e.g., row 1), Diet-Seg maintains segmentation consistency and avoids oversegmentation, likely due to its entropy-guided adaptive learning rate.

Overall, the qualitative results in Figure 5 confirm the superiority of Diet-Seg in both coarse tumor localization and fine-grained boundary precision. This is further supported by the clearer entropy maps and focused hardness prediction, which jointly highlight Diet-Seg’s ability to handle varying tumor size, morphology, and intensity contrast more effectively than the baseline Info Model.

To further investigate the pixel-level decision confidence of our model and the formation of the hardness map, we visualize the per-class entropy and its combinations in Figure 6. The first row presents: the input T1ce MRI image, the ground-truth annotation, and the prediction mask from our Diet-Seg model. The subsequent two rows display entropy maps for each tumor subregion (Class 1: WT, Class 2: TC, Class 4: ET), the entropy union of multiple subregions (Class 1+4, Class 1+2+4), and the final hardness map.

**Figure 6:**
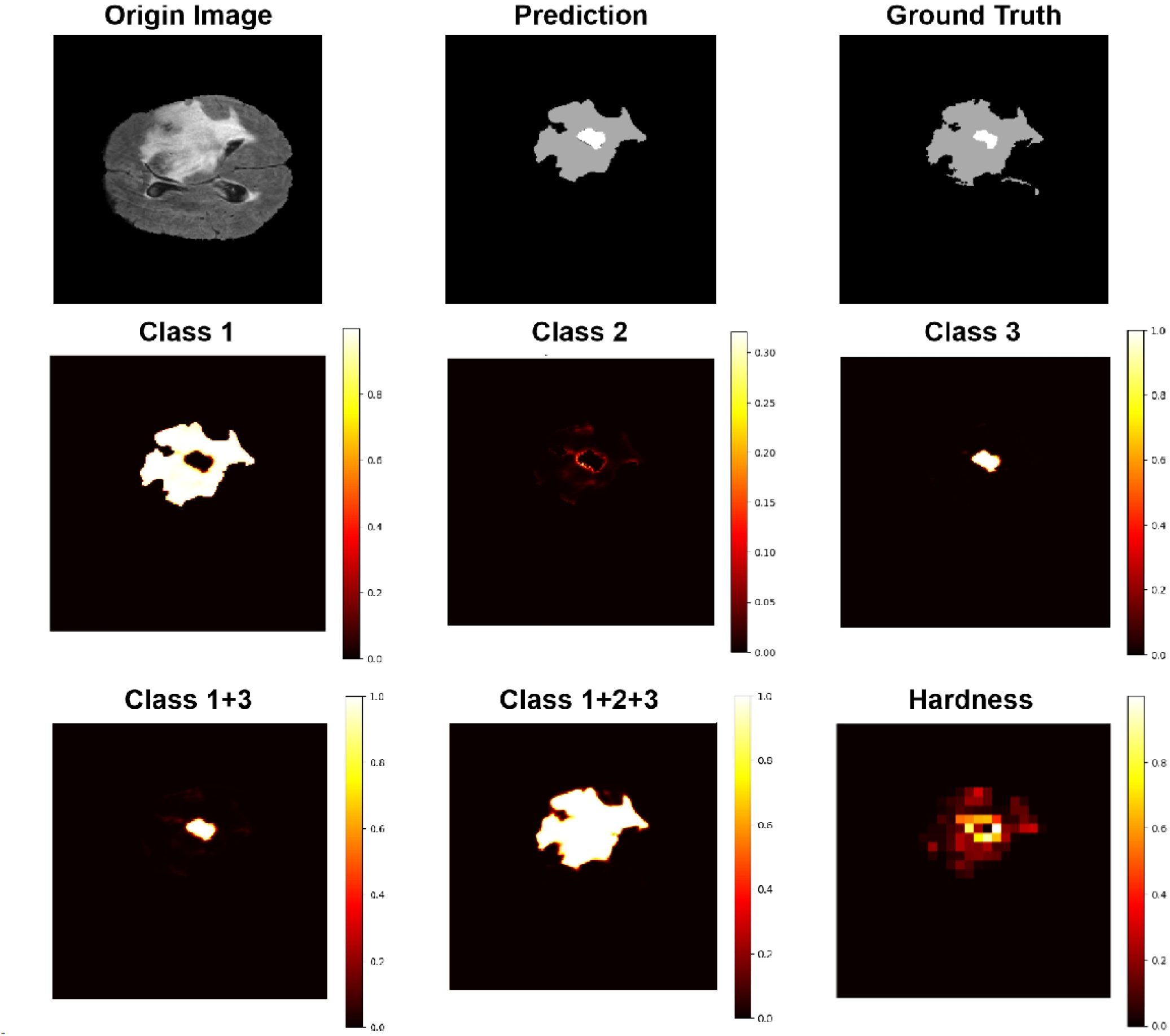
Entropy decomposition and hardness visualization.

As shown in Figure 6, the per-class entropy maps reveal distinct uncertainty patterns across different tumor subregions. The entropy for Class 1 (WT) tends to be higher across a broad area, reflecting the difficulty in delineating the entire tumor mass, especially near the edges. Class 2 (TC), which represents the tumor core, exhibits relatively lower entropy but with localized peaks around internal boundaries, indicating intra-tumor ambiguity. Class 4 (ET), which denotes the enhancing tumor, has highly concentrated uncertainty around small high-contrast areas, consistent with its anatomical appearance and clinical segmentation challenges.

The combined Class 1+4 and Class 1+2+4 entropy maps illustrate the aggregated uncertainty distributions. These merged entropy maps highlight areas with overlapping semantic ambiguity, emphasizing regions where tumor boundary definitions are difficult even for high-capacity models. Notably, the Class 1+2+3 map correlates well with the final hardness map on the right, demonstrating that our entropy-guided hardness computation effectively captures multi-class uncertainty and yields spatially meaningful training supervision signals.

Furthermore, the hardness map not only aligns with ambiguous boundary zones but also highlights regions of fine-grained complexity, such as fragmented enhancing tumors or low-contrast inner lesions. This supports the central hypothesis of our work: leveraging entropy-derived hardness maps allows the model to emphasize uncertain regions during training, resulting in more accurate and robust segmentation performance.

Overall, the decomposition and visualization validate the interpretability and effectiveness of the hardness-aware training mechanism within our Diet-Seg framework.

### 3.5 Ablation Study

To comprehensively evaluate the contribution of each component in our proposed architecture, we conducted an ablation study on the BraTS2019 dataset, as summarized in Table 4. The baseline model adopts a standard 3D U-Net trained as the information model, which is responsible for generating pixelwise hardness estimation and serves as the backbone foundation. Based on this, we incrementally added the RWKV-UNet module and EdgeNet component to observe performance changes in segmentation accuracy across tumor subregions: ET, TC, and WT.

**Table 4:** Ablation study of **EdgeNet, Info Model**, and **RWKV-UNet** on the BraTS2019 dataset. Dice scores are reported for each subregion.

| Configuration | ET (%) | TC (%) | WT (%) | Avg Dice (%) |
| --- | --- | --- | --- | --- |
| 3D U-Net (Info Model) | 81.33 | 82.54 | 79.50 | 81.12 |
| + RWKV-UNet | 85.92 | 86.24 | 83.71 | 85.29 (+1.17) |
| + EdgeNet (Full Diet-Seg) | **86.53** | **86.86** | **89.91** | **87.77** (+2.48) |

The 3D U-Net information model alone achieves Dice scores of 81.33% (ET), 82.54% (TC), and 79.50% (WT), yielding an average Dice score of 81.12%. This configuration serves as the reference point for evaluating architectural improvements.

Upon integrating the proposed RWKV-UNet module, which introduces long-range dependency modeling via recurrent weighted key-value interactions, the Dice scores improved substantially to 85.92% (ET), 86.24% (TC), and 83.71% (WT), with an average Dice of 85.29%. This corresponds to a significant

+4.17% gain over the baseline. The improvement verifies that the RWKV-based backbone effectively captures contextual and structural features across volumetric space, thereby enhancing segmentation robustness, especially in challenging regions like ET.

Further incorporating the EdgeNet module, which introduces edge-guided fusion and multi-scale attention to better preserve tumor boundaries, yields our final Diet-Seg configuration. With Dice scores of 86.53% (ET), 86.86% (TC), and 89.91% (WT), the full model achieves an average Dice of 87.77%, representing a +2.48% improvement over the RWKV-UNet alone and a +6.65% gain over the original 3D U-Net.

These results demonstrate that each architectural component in Diet-Seg provides meaningful and additive benefits. In particular, EdgeNet significantly boosts WT segmentation performance by enhancing spatial edge preservation, while RWKV-UNet enhances global context understanding for ET and TC. Together, they form a synergistic framework that achieves state-of-the-art segmentation accuracy in brain tumor analysis.

## Case Study

To further validate the clinical relevance of our proposed framework, we conducted a case study using a representative patient scan from the BraTS2019 dataset, characterized by irregular tumor morphology and low contrast boundaries. This case is visualized in Fig. 7.

**Figure 7:**
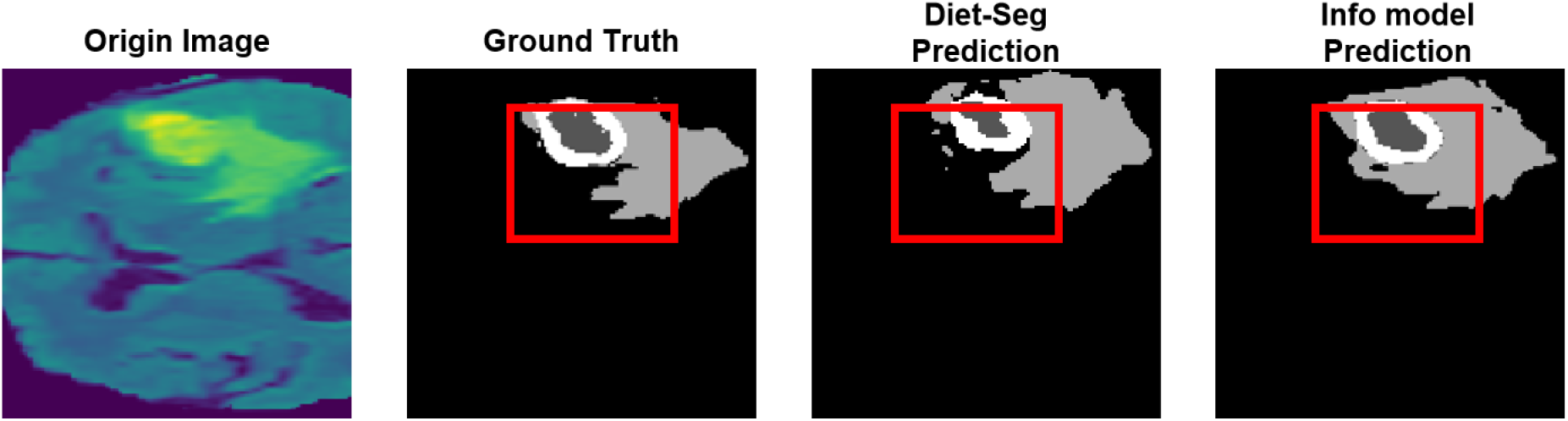
Case study on a BraTS2019 sample. From left to right: original MRI image (T1ce), ground truth segmentation, prediction by Diet-Seg, and prediction by the info model. Red box highlights the region where Diet-Seg outperforms in delineating tumor boundaries.

**Clinical Context** Accurate segmentation of ET, TC, and WT regions is critical for pre-operative planning and radiotherapy. This specific case involves a large glioblastoma with ambiguous ET boundaries and necrotic tissue in close proximity to the ventricles. Traditional models often over-smooth boundaries or misclassify adjacent edema and enhancing regions due to intensity similarity.

**Model Performance and Comparative Analysis** Figure 7 compares the ground truth annotations with the predictions of the baseline 3D U-Net information model and our Diet-Seg framework. As seen, the info model fails to clearly separate the enhancing tumor from surrounding edema, over-segmenting the WT region and leaking into normal tissue. In contrast, Diet-Seg provides a more faithful reproduction of the enhancing core and avoids false positives in the surrounding regions, indicating better spatial boundary preservation.

**Diagnostic Significance** From a clinical perspective, the accurate delineation provided by Diet-Seg holds significant implications. Over-segmentation, as seen in the info model, may lead to unnecessarily aggressive surgical margins or radiotherapy dosage beyond the tumor boundary, increasing the risk of healthy tissue damage. Conversely, Diet-Seg improves specificity by suppressing incorrect expansion and better respecting anatomical boundaries. This enhances safety during resection planning and supports more precise dose modulation in radiation oncology.

**Adaptivity via Hardness-Aware Training** The improved performance of Diet-Seg is attributed to its hardness-aware optimization strategy. During training, uncertain regions (such as the boundary between ET and TC) received adaptive learning rate adjustment based on entropy feedback from the info model. This allowed the network to pay more attention to hard-to-learn regions, mimicking the clinical workflow where radiologists iteratively focus on ambiguous zones.

This case highlights the strength of Diet-Seg in high-stakes clinical scenarios, where accurate localization of tumor boundaries directly informs therapeutic decisions. By decoupling the segmentation training from the info model, Diet-Seg achieves better adaptability and precision, suggesting strong potential for real-world deployment.

## Discussion

In this study, we proposed Diet-Seg, a novel training framework that integrates pixel-level hardness estimation and adaptive learning rate modulation to improve brain tumor segmentation. Extensive experiments demonstrate that Diet-Seg achieves consistent performance improvements across BraTS2018–2021 datasets, outperforming both traditional baselines and strong backbone models, including a high-performing 3D U-Net information model and RWKV-UNet. These gains validate the effectiveness of our entropy-guided dynamic optimization strategy in enhancing segmentation accuracy, particularly in anatomically ambiguous or low-contrast tumor regions. Compared to state-of-the-art models that rely on architectural innovations—such as advanced attention blocks, deeper networks, or multi-scale fusion—our contribution is fundamentally different. Diet-Seg does not introduce a new backbone, nor does it rely on feature-level fusion from auxiliary networks. Instead, it decouples the difficulty estimation process from the segmentation model by introducing a pretrained “information model” responsible for generating hardness maps that guide downstream optimization. This decoupling opens new possibilities for flexible, generalizable AI frameworks. The information model can be developed independently from the segmentation network, allowing clinical institutions to curate specialized hardness predictors tailored to specific imaging protocols or diagnostic tasks. This separation also aligns better with clinical learning processes, where radiologists iteratively adjust attention and confidence based on local ambiguity. Rather than injecting prior information into the input or feature space as in distillation or transfer learning, Diet-Seg adapts the learning behavior itself—a closer analogy to human expert learning. The resulting framework introduces a cognitively informed training paradigm, better suited to real-world diagnostic applications. From a deployment standpoint, Diet-Seg also offers practical advantages. The segmentation model trained under Diet-Seg retains only half the parameter volume of the original information model while achieving superior performance, making it more suitable for edge devices, mobile clinics, or real-time surgical navigation. Furthermore, its architecture-agnostic design allows seamless integration with a wide range of backbones, from UNet variants to Transformer-based architectures, broadening its applicability to diverse clinical settings and segmentation tasks (e.g., cardiac MRI, retinal vessel analysis, lung nodule detection). Nevertheless, our study has limitations. First, the additional computation required to generate hardness maps and apply adaptive optimization increases training time. Second, although we validate Diet-Seg across four datasets, further studies are needed to assess its robustness across institutions, scanner types, and disease categories. Lastly, the choice and quality of the information model directly affect hardness estimation accuracy, which in turn influences downstream performance. In future work, we aim to explore online updating strategies for information models, improve the sampling efficiency of hardness computation, and extend this framework to semi-supervised settings. Ultimately, Diet-Seg offers not just a performance boost but a new paradigm for learning in clinical AI—one that learns not only what to segment, but how difficult it is to learn.

## 4 Conclusion

We presented Diet-Seg, a novel brain tumor segmentation framework that dynamically modulates learning based on pixel-wise entropy-derived hardness. By leveraging a pretrained information model to estimate prediction uncertainty, Diet-Seg enables targeted optimization on challenging regions while preserving performance on easier structures. The integration of RWKV-UNet and EdgeNet further enhances long-range contextual modeling and edge detail preservation. Through extensive evaluation on the BraTS2019 and BraTS2020 datasets—as well as cross-year generalization on BraTS2018–2021—we demonstrate that Diet-Seg consistently outperforms existing state-of-the-art methods in both Dice Score and HD95 metrics.

Beyond tumor segmentation, the proposed hardness-aware learning paradigm offers a general and modular strategy that can be incorporated into various network architectures and extended to other medical imaging tasks. In future work, we plan to adapt this entropy-guided optimization to semisupervised learning, multi-modal segmentation, and large-scale clinical deployment scenarios. Overall, Diet-Seg establishes a strong foundation for developing adaptive, interpretable, and high-performance deep learning solutions in medical image analysis.

## Notes

### Competing Interest Statement

The authors have declared no competing interest.

### Funding

- National Health and Medical Research Council (NHMRC): 2041439

